# Conjunctive coding of letter pairs emerges in segregated human medial temporal lobe neurons during working memory

**DOI:** 10.64898/2026.06.04.729179

**Authors:** Esteban Félez Martínez, Filippo Costa, Debora Ledergerber, Lukas Imbach, Johannes Sarnthein, Timothée Proix

**Affiliations:** Institute of Neuroinformatics, University of Zürich and ETH Zürich, Zürich, Switzerland.; UniversitätSpital Zürich, Klinik für Neurochirurgie, Zürich, Switzerland.; Swiss Epilepsy Center, Klinik Lengg, Zürich, Switzerland.; Institute of Neuroscience, ETH D-HEST, Zürich, Switzerland.

**Keywords:** Single Unit Activity, Medial Temporal Lobe, Conjunctive Coding, Working Memory

## Abstract

Composing individual memory traces into unified representations is fundamental to encoding of structured relationships and flexible cognition. A central debate in neuroscience concerns the neural mechanisms of these compositions: are these compositions encoded through mixed selectivity, where the same neurons simultaneously represent individual and composed items, or through conjunctive coding within a distinct and specialized neuronal population? Here, we tested these competing hypotheses by recording 996 single units from the medial temporal lobe of 11 epilepsy patients performing a letter-string working memory task. We found neural encoding of both single letters and letter pairs during task related periods, with the latter peaking during the maintenance phase. Crucially, letter pairs were encoded by a sparse subset of neurons that did not show significant selectivity for their constituent letters. This suggests the existence of a segregated population of neurons recruited to compose items independently of single letter neural encoding. These findings provide evidence that the human medial temporal lobe separately encodes compositions within segregated neurons, offering direct neurophysiological support for conjunctive coding at the single-neuron level.

## 1 Introduction

In everyday cognition, we rarely process features in isolation; instead, we integrate items, contexts, and relations into structured, abstract representations that can be maintained, manipulated, and retrieved. This capacity underpins flexible cognitive behaviour across domains including perception, reasoning, language, and memory. Within the Language of Thought (LoT) framework, such abilities are taken to define the mind as a computational system that operates over symbolic representations and is inherently compositional and productive [1]. How such computational principles are implemented in neural substrates remains a matter of ongoing debate [2, 3].

In the recent years, the medial temporal lobe (MTL), classically known for its role in episodic memory and spatial navigation [4], has emerged as a potential key hub in the brain for the implementation of such symbolic representations and the computations over them [5–9]. A core function attributed to the hippocampal region of the MTL is the integration of distinct elements into structured relational representations that support episodic memory [10–13]. Growing evidence suggest that this integration is implemented through compositional coding, whereby neurons encode highly specific representations of particular combinations of variables or primitives (e.g., letter pairs) [14]. Studies in rodents have demonstrated that single neurons can encode these combinations of variables, for instance in the head direction or spatial location systems [15]. Recent work suggests that the hippocampus supports compositional coding by combining reusable primitives into structured representations, highlighting the role of hippocampal replay to generalize and infer novel states, thereby enabling flexible cognition beyond episodic memory and spatial navigation [16]. Complementary theoretical and empirical work indicates that these representations may be implemented in population activity, where separate populations represent single and composed elements, offering a mechanism for the stability and flexibility of compositional codes in the MTL [**?**].

Despite these theoretical advances, the neuronal basis of compositional representations remains poorly understood. In humans, prior work in the MTL has largely focused on selectivity for individual items, including concept cells that respond to the abstract identity of a specific individuals [17, 18], as well as neurons tuned to single symbolic elements [19–22]. However, the extent to which single neurons encode combinations of items beyond the individual primitives, and whether such selectivity depends on task context, has not been systematically examined. Two competing hypotheses are possible (Fig. 1). Encoding compositions could be performed through conjunctive coding (H1), where a dedicated subset of neurons that selectively encode specific item combinations [23]. Alternatively, compositions may arise through mixed coding (H2), in which individual neurons jointly encode both constituent primitives and their conjunctions [24, 25]. Both of these forms of multi-dimensional encoding could provide the flexibility required to represent complex relationships while preserving the specificity necessary for episodic memory.

**Fig. 1:**
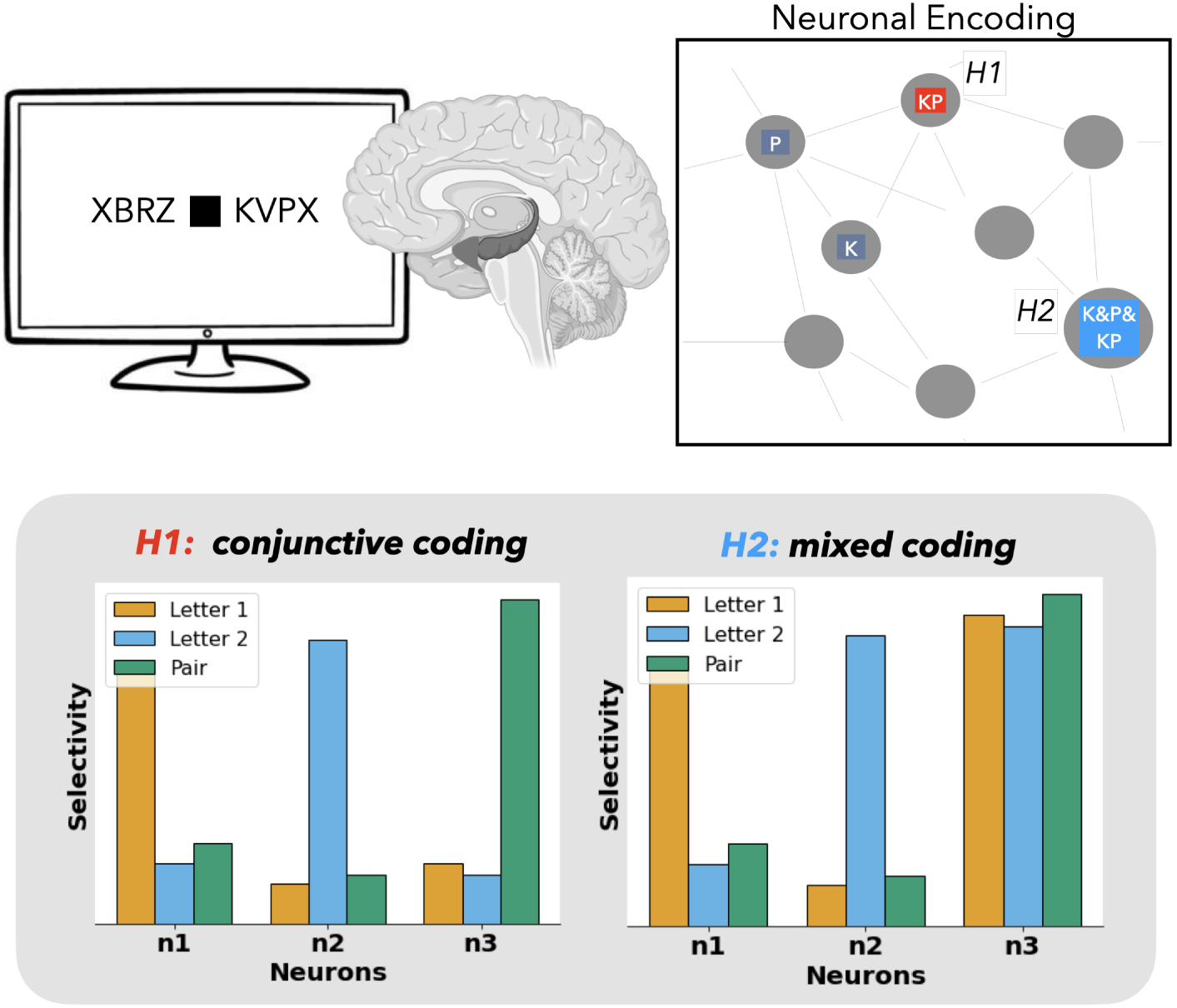
Conjunctive coding (H1) versus mixed coding (H2) hypotheses. We recorded single-neuron activity in the medial temporal lobe of epilepsy patients during an experiment which involved keeping a letter string in working memory. Two competing hypotheses for how neurons encode letter pairs were tested: H1 (conjunctive coding) predicts that neurons encode letter pairs independently of individual letter encoding. H2 (mixed coding) predicts that neurons encoding letter pairs also encode individual constituent letters.

Here, we contrasted these two hypotheses experimentally by recording individual neurons (n = 996) recorded from the MTL of 11 epilepsy patients during a modified Sternberg task involving keeping a letter string in working memory [26, 27]. Using generalized linear models and an information-theoretic measure of selectivity, we identified neuronal responses that jointly coded for letter pairs (e.g., KP), beyond the independent contributions of their constituent letters (e.g., K). We further examined whether single letter and letter-pair representations co-occurred within the same neurons, or were segregated across distinct neuronal populations (Fig. 1). We found that letter-pair coding emerges in a subset of neurons varying across task epochs, with strongest effects during the memory maintenance period of the task. Notably, bigram coding was predominantly observed in neurons that did not exhibit significant single-letter selectivity under our model-based criterion, suggesting partial segregation from neurons showing single-letter selectivity, consistent with the pure conjunctive coding hypothesis (H1). Overall, these results indicate that the human MTL supports both independent and conjunctive representations of task-relevant items during working memory.

## 2 Results

### 2.1 Task and recordings

We recorded single-neuron activity of eleven epilepsy patients performing a modified Sternberg working memory (WM) task (Fig. 2A) [5, 27]. Each trial began with a Fixation stage of 1 second, followed by the presentation of a set of 4, 6 or 8 letters (Encoding, 2 s) drawn from a pool of 15 consonants. After a Maintenance stage (3s), a probe letter was presented and participants had to retrieve from memory (Retrieval, 2 s max) and to indicate by button press (“IN” or “OUT”) whether the probe letter was a member of the letter set. During the Maintenance period, participants rehearsed the phonemes of the letter strings subvocally, as instructed, and confirmed using this strategy after the sessions. This activation of the phonological loop is a component of verbal WM supporting appropriate behavioral responses [28, 29]. Unlike the original Sternberg’s paradigm, all items were presented simultaneously rather than sequentially.

**Fig. 2:**
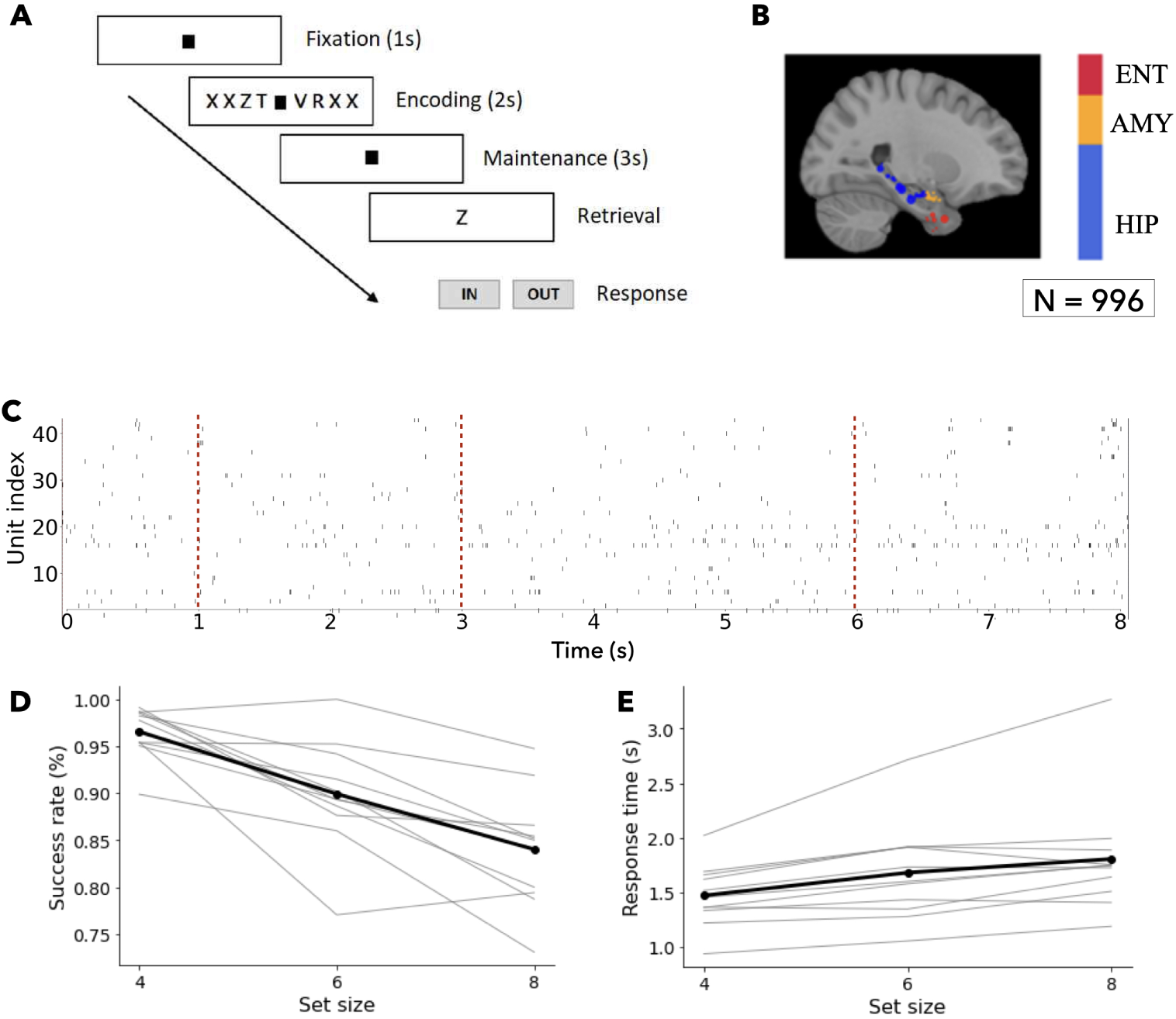
Single-neuron recordings from 11 epilepsy patients during a modified Sternberg’s task with letter strings. **A** The modified Sternberg task involves remembering whether a probe letter was part of a presented letter strings. **B** We recorded single-unit data in 11 subjects from the hippocampus (HIP), amygdala (AMY) and entorhinal cortex (ENT), yielding a total of 996 units. **C** Raster plot for the recorded single-units during one trial for one of the patients. Vertical lines identify trial period transitions. **D-E** Success rate decreased and response time increased with set size (grey: individual patients; black: mean)

Single-neuron activity was recorded via depth electrodes implanted in the medial temporal lobe (Fig. 2B). We identified a total of 996 putative single units (hereafter referred to as ‘neurons’) across all microelectrodes: 559 in the hippocampus (HIP, 56%), 239 in the amygdala (AMY, 24%), and 198 in the entorhinal cortex (ENT, 20%). An exemplary raster plot of the single-neuron data is shown in Fig. 2C.

Behavioral performance was high, with average correct response rates of 92 ± 5.3% for IN trials and 91 ± 4.0% for OUT trials. The correct response rate decreased with set size (97% for set size 4, 90 % for set size 6 and 84 % for set size 8, permuted repeated-measures ANOVA F(2, 16) = 32.27, *p <* 0.0001, Figure 2D). Across all participants, the capacity averaged 5.52 [Cowan’s K, (correct IN rate + correct OUT rate −1) x set size]. Correct IN/OUT decisions were made more rapidly than incorrect decisions (1.56 ± 0.37 s vs. 1.89 ± 0.57 s, permutation t test, *p* = 0.0020). Consistent with Sternberg’s scanning hypothesis that the participants kept a stable representation of the stimulus in WM [30], response time increased with set size (permuted repeated-measures ANOVA F(2,16) = 14.25, *p <* 0.0001, Fig. 2E) by 63 ms/item. The distribution of letter occurrence, recency, and primacy in the Encoding set did not significantly differ from a uniform distribution (chi2 test *p >* 0.14).

### 2.2 Single-letter encoding peaks during task engagement

We first asked whether individual neurons encoded the identity of single letters, independently of other letters. To do so, for each neuron, letter, and task period, we compared two point-process generalized linear models (PPGLMs) of spiking activity: a null model containing task-related confounders (trial set size, trial period, and single-neuron spike history), and a full model that additionally included letter identity (Fig. 3A). The increase in model performance was quantified in bits per spike (BPS), providing an information-theoretic estimate of the improvement in spike response prediction brought by letter identity [31]. In practice, this measures how much more accurately the full model predicts the timing and probability of neuronal spikes when information about letter identity is included. The BPS value can be interpreted as a gain in predictability obtained by adding the new covariates to the model. In Fig. 3B, we show, for each letter and trial period, the maximum BPS across all the tested neuronal models (i.e., across all neurons). Single-letter encoding was weakest during Fixation, which marks the noise baseline, and substantially stronger during the task-related periods. The total information carried by the population peaked during Maintenance and remained elevated during the Retrieval period, indicating that letter-specific representations were strongest once the stimulus had been encoded and had to be actively maintained or retrieved.

**Fig. 3:**
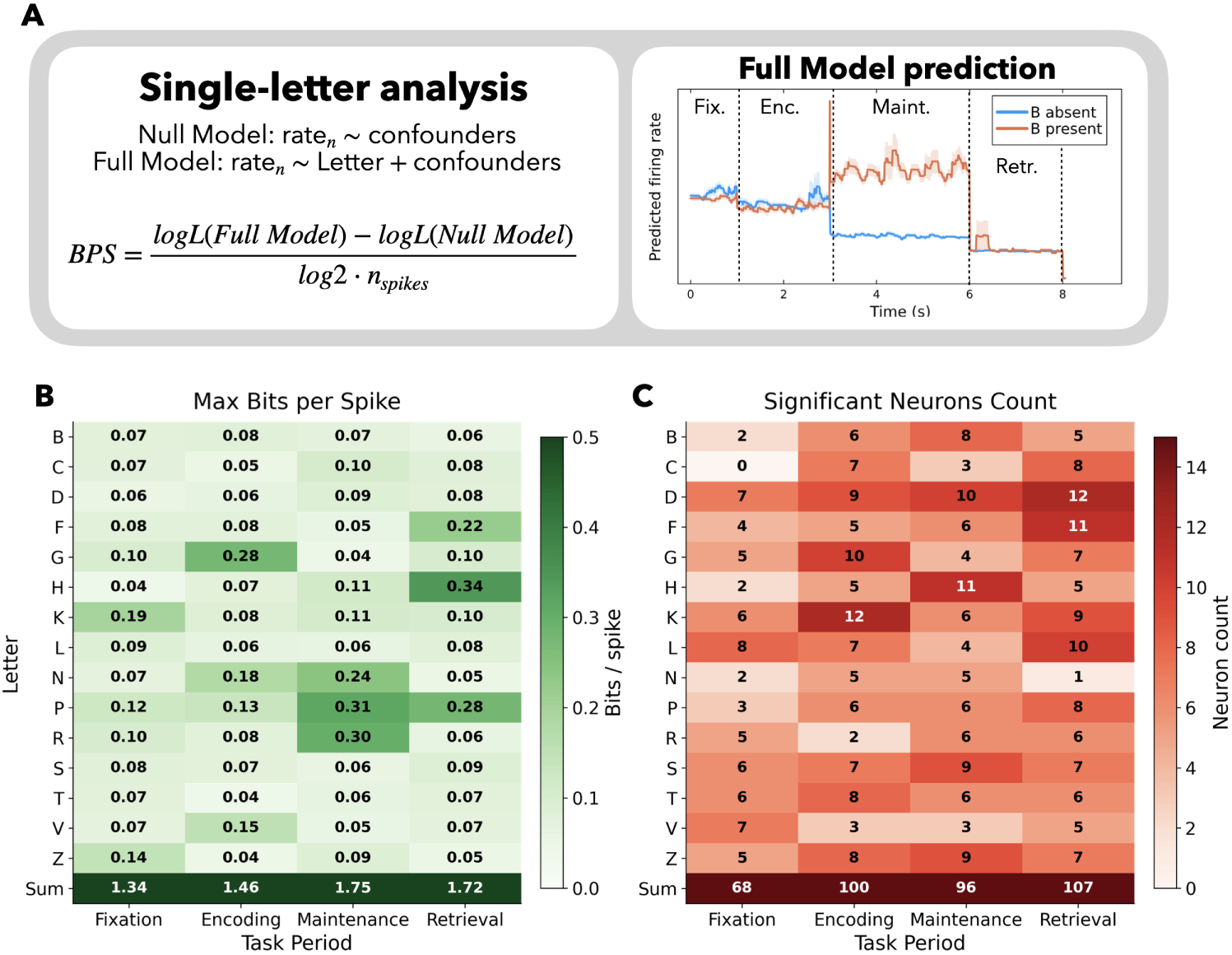
Maximum single-letter encoding across neurons during task-related periods. **A** Single-letter analysis approach. Left: the full model includes Letter identity as a predictor alongside confounders, compared to a null model with confounders only. Bits per spike (BPS) is used to quantify the information gain provided by the full model with respect to the null model [31]. Confounders include trial period, spike history and trial set size. For a single session, all trials were concatenated and the model was run independently for every neuron, letter and trial period. Right: Full model prediction for a neuron encoding letter B during Maintenance. **B** Heatmap of maximum BPS across neuron-specific models for each letter identity across the four task periods. Values represent the highest BPS for each letter within each task period across all neurons and patients. The bottom row (Sum) shows the total information across all letters per period, with Maintenance and Retrieval periods showing the highest overall encoding. **C** Significant number of neurons encoding single letters. This number is significantly greater during Encoding, Maintenance and Retrieval periods compared to Fixation (permutation test, *p <* 0.05). Encoding neurons are found using a permutation test at (*α* = 0.05, FDR-corrected), in which we run the model n = 200 times randomizing the letter appearance.

We assessed the significance of each neuron’s BPS using a null distribution generated by randomly permuting letter labels (see Methods). Briefly, neurons were classified as significantly encoding if they passed a permutation test (*α* = 0.05, FDR-corrected), in which the model was rerun n = 200 times after randomizing letter appearance labels. In Fig. 3C, the total number of neurons passing the test for each letter and trial period is shown. The total number of neurons that passed the permutation-based single-letter selectivity test was minimal during Fixation and significantly greater during Encoding, Maintenance, and Retrieval (*p <* 0.05, permutation test). We also examined whether letter-encoding neurons are shared across trial periods by assessing the overlap of encoders across the four phases (Fig. A1). Most neurons encode letters in a period-specific manner, with limited overlap observed primarily between the Encoding-Maintenance and Retrieval-Maintenance periods. This analysis revealed clear letter-selective responses across the population, with a marked specificity for each task epoch.

### 2.3 Letter-pair encoding shows strongest signal during Maintenance

We next asked whether neuronal responses carried information about specific combinations of non necessarily adjacent letters (called letter pairs thereafter) beyond the contribution of the individual letters themselves. For each neuron, letter pair, and task period, we therefore compared a null model containing the identities of the two constituent letters and the same task-related confounders as above, with a full model that additionally included their interactions (Fig. 4A). This model is intended to capture letter pair compositional representations without any assumptions about single letter encoding. In this analysis, a positive increase in BPS indicates that a model including pair identities explains spiking activity better than a model without it, thereby isolating composition effects. Similar to the previous analysis, significance was assessed using a null distribution generated by randomly permuting letter-pair labels. In this case, the permutation test revealed no significant differences in the number of neurons encoding letter pairs across trial periods.

**Fig. 4:**
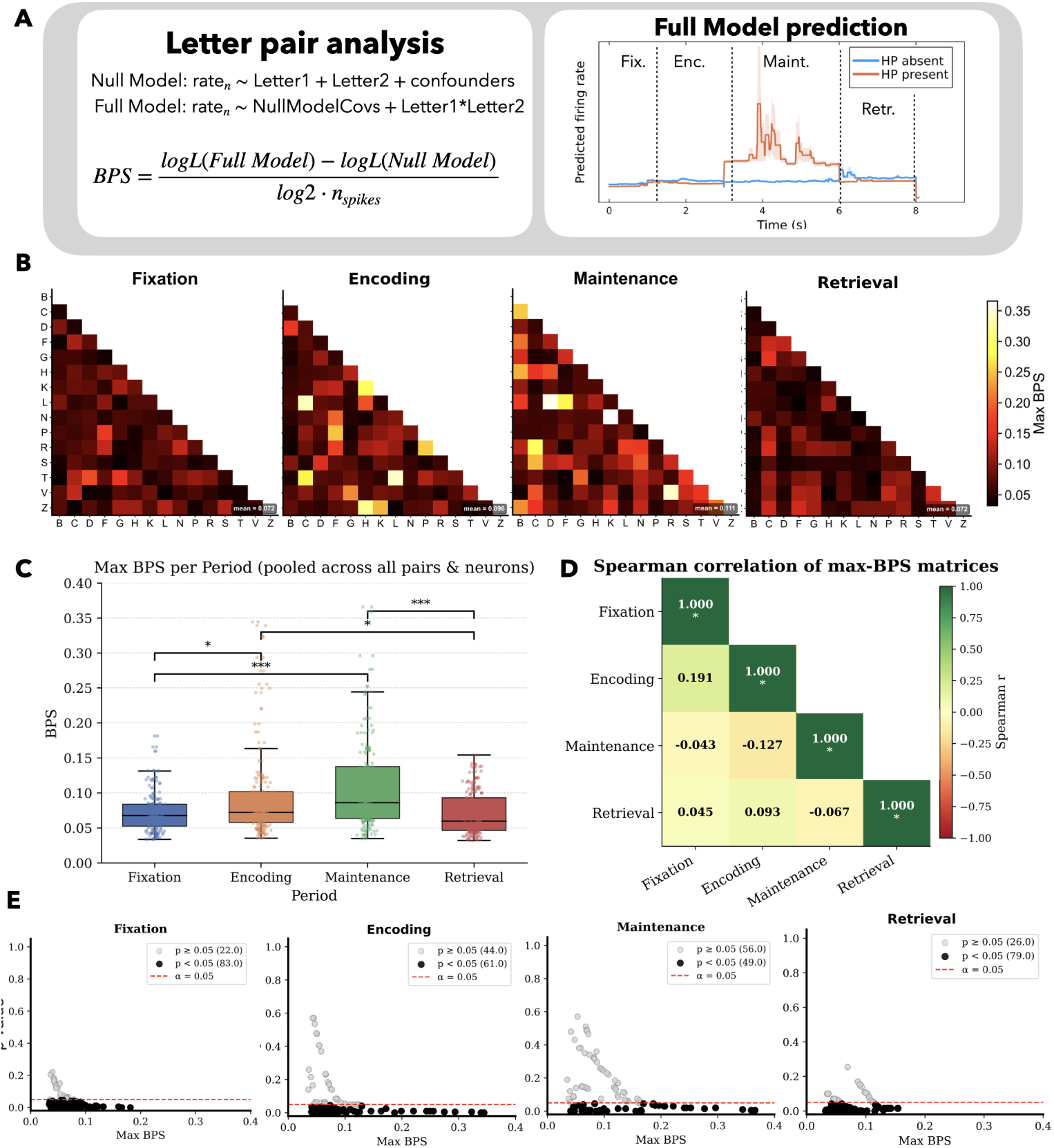
Letter pairs are best encoded during the Maintenance period. **A** Letter-pair analysis approach. The null model includes two Letter identities as predictors alongside confounders, while the full model adds the interaction between such letters. Confounders include trial period, spike history and trial set size. For a single session, all trials were concatenated and the model was run independently for every neuron, letter pair and trial period. The schematic in the right shows the difference between the Full and Null model predictors. Right: Full model prediction for a neuron encoding letter pair HP during Maintenance. **B** Maximum BPS value across all 996 neuronal models for each letter pair, shown separately for each trial period. Rows and columns represent letters, and each matrix entry corresponds to the maximum BPS across neurons for the given letter pair. **C** Distribution of the maximum BPS values shown in B across trial periods. Boxplots indicate that peak encoding strength was highest during the Maintenance period. Statistical significance was assessed using Wilcoxon tests (∗*p <* 0.05, ∗ ∗ ∗*p <* 0.001*, BC*). **D** Spearman’s rank correlation across the four upper-triangular maximum-BPS matrices (B). Correlations with the Maintenance period are all negative. **E** For all the neuronal models that gave the max BPS results in B, we show the p-value of the corresponding test (permutation test shuffling letter-pair labels).

To further evaluate letter-pair encoding strength beyond individually significant neurons, we examined the maximum BPS value obtained across the 996 neuronal models for each letter pair in each trial period (Fig. 4B). Across periods, the highest maximum BPS values were concentrated during the Maintenance epoch, indicating stronger pair-specific encoding during this stage of the task. To summarize these differences, we compared the distributions of maximum BPS values across periods (Fig. 4C). Maximum BPS values were significantly greater during Maintenance than in Fixation and Retrieval (*p <* 0.001, Wilcoxon tests, Bonferroni-corrected), and greater during Encoding compared to Fixation and Retrieval (*p <* 0.05, Wilcoxon tests, Bonferroni-corrected). In further analysis, we focus on the Maintenance period, as this phase is the most likely to reflect the emergence of letter-pair representations. The non-zero results for Fixation can be interpreted as a chance-level baseline, as no letter information was present in this stage.

In addition, the structure of pair selectivity differed across epochs: correlations between the upper-triangular entries of the maximum-BPS matrices were negative for the Maintenance period relative to the other periods (Fig. 4D), suggesting that the letter-pair combinations most strongly represented during Maintenance were different from those already encoded during Fixation (which captures baseline variability), Encoding or Retrieval periods. Finally, we asked whether the neuronal models underlying the maximum BPS cases were significant under our letter-pair permutation test (Methods), or were just a mere consequence of some task unrelated neural features. In Fig. 4E, we show that most of the neuronal models (all above the 0.2 BPS threshold) driving the maximum-BPS results are also significant according to the permutation test on letter-pair labels.

Together, these results indicate that compositional representations of letter pairs are enhanced during memory maintenance and cannot be explained by a static pattern of elevated baseline selectivity present across trials, as reflected in the negative correlation observed in the Maintenance period (Fig. 4D). This interpretation is further supported by the low overlap in the neurons showing maximal letter-pair BPS across task periods (Supplementary Fig. A2), indicating that letter-pair selectivity is not consistently driven by a fixed subset of neurons. Instead, these findings suggest that compositional selectivity emerges in a dynamic and task-dependent manner across the population.

### 2.4 Maintenance-specific compositional neurons encode letter pairs

To determine whether the Maintenance-related composition signal reflected a broad population trend or a sparse subset of highly selective neurons, we next examined each neuron’s maximal selectivity across letter pairs during Maintenance relative to its baseline pair-selectivity during Fixation. Across neurons, maximum BPS values during Maintenance were overall positively related to those observed during Fixation, Encoding, and Retrieval (Fig. 5A), indicating that some neurons showed generally higher composition-related model gains than others, possibly due to specific firing rate patterns. We therefore used a regression-based approach to identify neurons whose Maintenance-period compositional encoding exceeded the level expected from their Fixation baseline. Specifically, we regressed each neuron’s maximum Maintenance BPS across letter pairs onto its maximum letter-pair Fixation BPS, and identified the neurons with residuals greater than two standard deviations above the regression fit as Maintenance-specific compositional encoders (Fig. 5B).

**Fig. 5:**
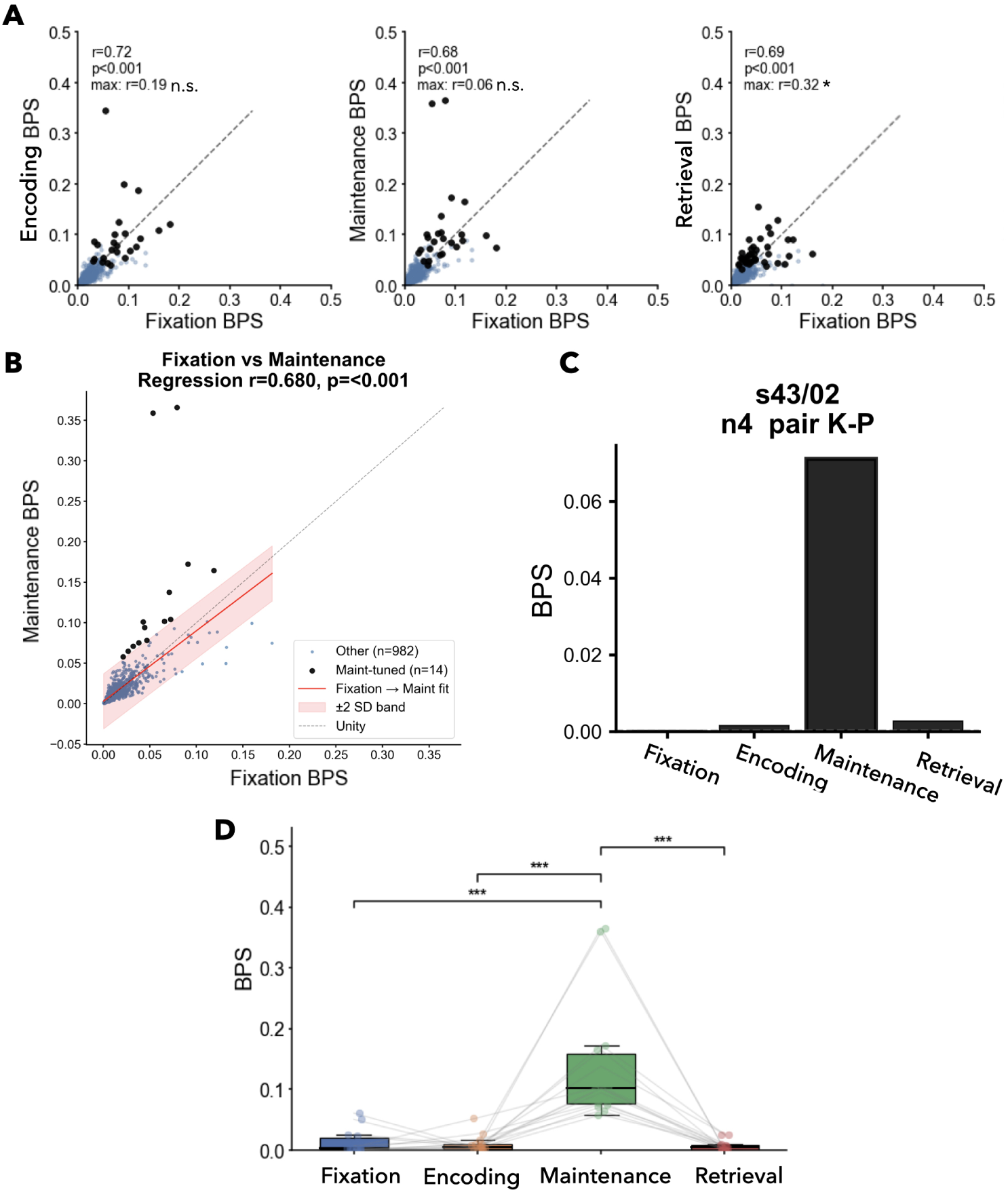
Letter-pair encoding neurons during Maintenance period. **A** Scatter plot of per-neuron maximum BPS across letter pairs during Fixation (x-axis) versus Encoding, Maintenance and Retrieval (y-axis), where each point represents one neuronal model (n=996). Black dots identify the neuronal models driving the results shown in Fig. 4B. Pearson’s correlation was used to quantify the linear relationship between each neuron’s maximum BPS during periods across all letter pairs. **B** We regressed each neuron’s maximum BPS during Maintenance onto its maximum BPS during Fixation to identify Maintenance-specific compositional neurons. These were identifed as neurons whose Maintenance BPS exceeded the regression-predicted value by more than 2 standard deviations of the residual distribution. 14 neurons passed this test. **C** BPS during the four task periods of an example of one of the 14 Maintenance-tuned neurons (in this case, K-P). **D** BPS across the 14 Maintenance-tuned neurons for the four task periods. The same letter pair is tracked across all four periods for each neuron, such that the BPS values reflect the coding of a specific letter pair rather than the pair maximized for each period independently. Individual data points and their trajectories across periods (grey lines) are shown for each neuron. Brackets indicate pairwise Wilcoxon one-sided signed-rank tests comparing each period against Maintenance. Significance levels: ∗*p <* 0.05, ∗ ∗ *p <* 0.01, ∗ ∗ ∗*p <* 0.001.

Using this criterion, we identified a sparse subset of 14 neurons whose letter-pair encoding was selectively enhanced during Maintenance (Supplementary Fig. A3A). These neurons come from four of the eleven patients, and from the anterior hippocampus (1 neuron), posterior hippocampus (9 neurons), amygdala (2 neurons) and entorhinal cortex (2 neurons). Fig. 5C illustrate an example neuron with strong pair-specific coding for a particular letter-pair during the Maintenance period. When the same letter pair was tracked across all task periods for these 14 neurons, pair coding was highest during Maintenance and significantly lower in Fixation, Encoding, and Retrieval (one-sided Wilcoxon signed-rank tests; Fig. 5D), confirming that these cells were tuned to pair identity specifically while the letters had to be held in memory. Thus, although the identified subset is small, our analysis isolates a group of neurons with Maintenance-enhanced conjunctive responses that are not explained by baseline pair selectivity. Importantly, these findings did not survive after two controls: when regressing Encoding to Fixation BPS to find Encoding-specific coders (Supplementary Fig. A3B,C) and when inverting the baseline regression to find Fixation-specific coders compared to a Maintenance baseline (Supplementary Fig. A3D).

### 2.5 Conjunctive coding in compositional neurons: lack of encoding for constituent letters

We finally asked whether neurons showing Maintenance-specific letter-pair coding also encoded the individual letters composing those pairs (mixed coding), or constituted a distinct set of selective letter-pair encoding neurons (conjunctive coding) (Fig. 1). For each of the 14 Maintenance-tuned letter-pair encoding neurons, we examined the single-letter BPS values associated with the two constituent letters of the preferred pair across all task periods. In Fig. 6A, several representative example neurons are shown, in which robust Maintenance-period pair coding occurred despite weak or absent single-letter selectivity for the constituent letters.

**Fig. 6:**
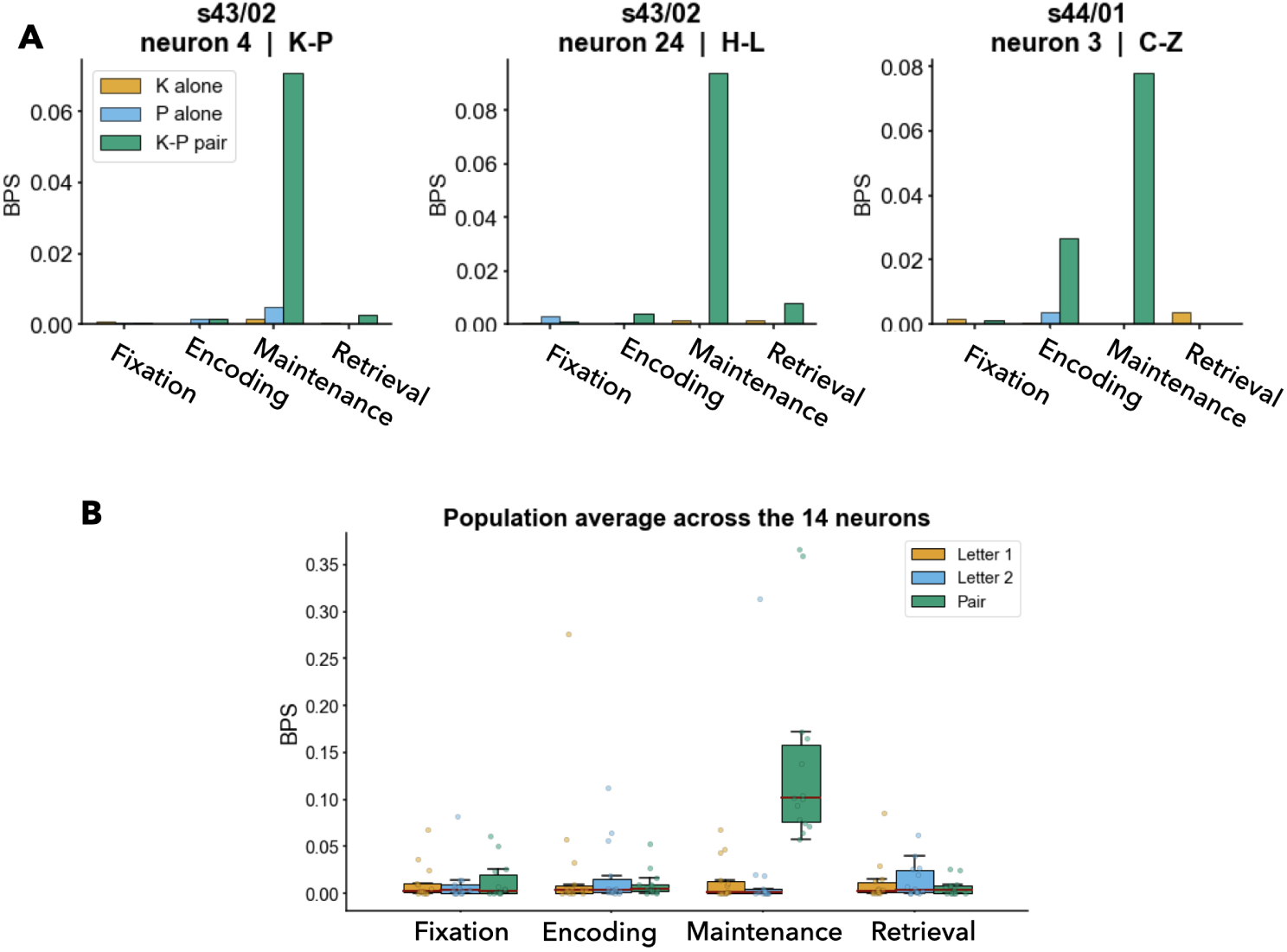
Conjunctive coding of letter pairs: compositional neurons do not encode constituent letters. **A** BPS of single constituent letter and letter-pair models across periods for three example neurons. **B** Selectivity results across the 14 letter-pair encoders.

This pattern was maintained at the group level. Averaging across the 14 letter-pair encoding neurons showed that single-letter selectivity for the letters composing the preferred pair remained modest across task periods, particularly relative to the strong Maintenance-period pair signal that defined these neurons (Fig. 6B), consistent with our conjunctive coding hypothesis (H1, Fig. 1). Importantly, we ran a control analysis in which we selected the 14 neurons with the highest single-letter selectivity during the Maintenance period, independently of any pair-based criterion. For each of these neurons, we identified the two most strongly encoded letters and compared their responses to the maximal pair response across all possible letter combinations. Results, shown in Supplementary Fig. A4, demonstrate that pair-related BPS remained comparatively weak, indicating that strong single-letter tuning does not give rise to pair encoding.

## 3 Discussion

In this study, we aimed to to test whether single neurons in the human medial temporal lobe encode specific combinations of symbolic items during working memory. Across 996 neurons recorded from 11 patients during a modified Sternberg’s task involving letters, we found that letter-pair coding, quantified by the model-based measure of BPS, was strongest during the Maintenance period and that the neurons with the clearest maintenance-related pair coding generally showed weak selectivity for the constituent letters of their preferred pair. Taken together, these results suggest that the human MTL contains a sparse conjunctive code for item combinations that becomes prominent when information must be actively maintained.

Our findings extend previous work showing that neurons in the human MTL can be selective for individual concepts or symbolic elements [17–22]. Here, the critical result is not simply that neurons responded during a verbal memory task, but that for some neurons the response to a letter pair could not be predicted from the additive contribution of the two letters alone. This is consistent with proposals that the MTL supports compositional operations that integrate multiple elements into structured representations [11–13]. Our data suggests that conjunctive coding may contribute not only to long-term relational memory, but also to the active maintenance of items during working memory. Whether the same units dynamically appear temporarily to solve the task remains an open question, requiring experimental designs with progression across condition and demand. [32].

Importantly, our findings were strongly dependent on the trial period. Single-letter coding increased during the task-relevant periods, but conjunctive coding peaked specifically during Maintenance, when the stimulus was no longer present and had to be kept internally. This argues against a purely sensory explanation of pair coding and instead supports the view that conjunctive representations are recruited when the task requires a bound internal representation of the memorized stimulus. Such flexibility in adaptation to task demands has been previously reported in MTL neurons [33].

A central question in this study was whether compositional coding co-occurs with single-letter coding in the same neurons (H2: mixed coding) or instead emerges in a partially distinct subpopulation (H1: conjunctive coding). Our results are more consistent with the latter interpretation. Among the neurons with maintenance-enhanced pair coding, selectivity for the constituent letters was generally modest, indicating that pair coding need not arise as a simple consequence of strong tuning to the individual letters. This does not imply a complete separation between single-item and conjunctive coding across the MTL; rather, it suggests that multiple coding schemes may coexist within the same system, with some neurons preferentially tracking individual primitives and others preferentially tracking their conjunctions [23]. Such a coding regime may be particularly useful for memory systems, where representing specific combination of items in specialized neural subsets might be key to create structured memories [21].

The exact operations performed by the neuronal circuits to perform such conjunctive coding can only be hypothesized. The finding that pair coding peaks during maintenance suggests an active integration of information units, rather than a purely stimulus-driven response since the letters were no longer visually present during this phase, which aligns with the proposal that conjunctive representations are stored in hippocampal memory through inter-hippocampal connectivity [16]. In this regard, one possible mechanistic interpretation is that such conjunctive signals arise from multiplicative interactions between single-letter representations, effectively implementing a form of outer-product coding between neuronal populations [34]. However, this type of explicit combinatorial expansion scales poorly: the number of possible pairwise combinations grows quadratically (*n*^2^) with the number of primitives (*n*), with higher-order conjunctions increasing even more rapidly. This combinatorial explosion makes a fully conjunctive scheme difficult to maintain as a general-purpose coding strategy beyond relatively small and structured feature spaces, suggesting that conjunctive coding may be most plausible for low-complexity representational regimes (such as letter pairs, syllables, or other constrained sets of frequently co-occurring items like words) while more distributed or compositional mechanisms are likely required as representational demands increase [35]. This trade-off between multi-item conjunctive and distributed representations could explain the compositional and productive capacity of the human mind, as we can easily depict the composition of concepts *a priori* unrelated, such as a flying red horse [1–3]. In this sense, the observed pair coding in MTL may reflect a regime in which explicit conjunctive representations are still tractable and useful, rather than a general solution for arbitrarily complex structured memory.

There are some limitations to our work. First, the Maintenance-specific subset of pair coding neurons was small, comprising 14 neurons selected based on the Maintenance-Fixation regression analysis. As a result, some neurons representing letter pairs may not have met the criteria for inclusion in this test. Second, the recordings were obtained from epilepsy patients implanted for clinical purposes, and sampling across MTL subregions was constrained by clinical needs rather than experimental design. It should also be noted that epilepsy-related activity, such as interictal spikes or ripples, may interfere with the identification of single-neuron activity. Third, our stimuli were limited to letter strings in a verbal working-memory task, so it remains unclear whether the same coding principle generalizes to nonverbal stimuli, more abstract relations, or episodic memory paradigms. Then, the possibility of a sampling bias cannot be excluded, which may have resulted in the omission of H2 neurons. Finally, our letter-pair modelling framework did not account for letter positions in the presented string, treating cases where the letters appeared together as indistinguishable from those where they were spatially separated. A natural extension of this work is to overcome these limitations by using data from more patients, applying our framework to non-verbal (and potentially non-visual) stimuli in a similar task structure, and employing more exhaustive models that account for item order.

In conclusion, our findings suggest that the human MTL encodes not only individual items but also specific item combinations during working memory in a specialized manner, consistent with our proposed conjunctive coding hypothesis and some of the findings in the literature [23]. By demonstrating that cognition relies on operations performed over symbolic and discrete representations, this study lends empirical weight to the LoT framework [1], pointing towards the MTL (and the single neurons therein) as a key substrate for these integration operations.

## 4 Methods

### 4.1 Dataset

Eleven patients with epilepsy implanted with depth electrodes in the MTL for the potential surgical treatment of epilepsy participated in the study. Placement of the electrodes was only guided by clinical reasons related to epilepsy. All participants had normal or corrected-to-normal vision and were right handed, as confirmed by neuropsychological testing. All participants provided written informed consent for the study, which was approved by the institutional ethics review board (PB 2016-02055). We used a modified version of the traditional Sternberg task [30], in which the encoding of memory items, maintenance, and recall were temporally separated (Fig. 2A) [5]. Each trial started with a fixation period of 1 second, followed by a 2-second presentation of the stimulus, which consisted of a set of eight consonants at the center of the screen. The middle four, six or eight letters were the memory items, which determined the set size for the trial (4, 6 or 8, respectively). The outer positions were filled with “X”, which was never a memory item. After the stimulus period, the letters disappeared from the screen, and a 3-seconds maintenance interval started. A fixation square was shown throughout fixation, encoding, and maintenance. After maintenance, a probe was presented in the response period. The participants responded, as rapidly as possible and without making errors, with a button press to indicate whether or not the probe was part of the stimulus (”IN” vs “OUT”). After the response, the probe was turned off, and the participants received acoustic feedback regarding whether their response was correct or incorrect. During the maintenance period, participants were instructed to rehearse the phonemes of the letter strings subvocally, and they confirmed using this strategy after the sessions. This activation of the phonological loop is a component of verbal WM supporting appropriate behavioral responses [28, 29].

Importantly, the distribution of letter occurrence, recency, or primacy in the stimulus set did not significantly differ from a uniform distribution, therefore, results are not driven by biased letter presentation numbers.

### 4.2 Recording setup and unit identification

Depth electrodes (1.3 mm diameter; 8 contacts of 1.6 mm length; 5 mm spacing between contact centers; ADTech®, Racine, WI, USA) were stereotactically implanted in the amygdala, hippocampus, and entorhinal cortex. Each macroelectrode contained nine microelectrodes extending approximately 4 mm beyond the electrode tip. Neural signals were acquired using the ATLAS recording system (Neuralynx, Bozeman, MT, USA) at a sampling frequency of 32 kHz with a 0.5–5000 Hz passband. Spike sorting was performed using the Combinato package [36]. Similar to other commonly used spike-sorting approaches, Combinato detects peaks in the high-pass filtered signal (*>*500 Hz), extracts wavelet coefficients from detected events, and applies superparamagnetic clustering within the resulting feature space. Compared with alternative clustering methods, Combinato provides improved automated artifact rejection and enhanced sensitivity for detecting small clusters containing relatively few action potentials.

All identified clusters were visually inspected based on waveform shape and amplitude, as well as interspike interval (ISI) distributions. Each candidate unit was passed through an automated stage and rejected if it failed any of the following criteria: a mean waveform peak exceeding +150 µV or a trough below 150 µV; a refractory-period contamination above 4%, defined as the fraction of inter-spike intervals 3 ms; a median peak-to-peak amplitude below 40 µV; an absence of sustained activity across the recording; a waveform-shape constraint requiring physiological polarity between the early and late portions of the mean waveform. Units were additionally rejected if their mean firing rate fell below a minimum threshold. This threshold was adjusted on a per-session basis rather than held fixed, in order to accommodate recordings with sparse spiking. Units surviving automated rejection were subsequently curated manually. During this step we primarily assessed the shape of the mean waveform and the presence of persistent spiking throughout the recording. As a final step, clusters belonging to the same micro-contact were inspected using their overlaid mean waveforms and their peak-to-peak amplitude over time, and were either merged or left separate to yield the final set of single units. This yield a total number of 996 putative single units (neurons) across the 11 participants: 559 units in the hippocampus (HIP, 56%), 239 units in the amygdala (AMY, 24%), and 198 units in the entorhinal cortex (ENT, 20%) (Fig. 2B).

### 4.3 Single-letter analysis

To identify neurons encoding single-letters in specific task periods, we used a point-process generalized linear model (PPGLM) approach. First, spike times were binned (width of 10 ms) and trials in a given session were concatenated, using a one-second intertrial intervals to avoid cross-trial interference. Then, for every neuron *n*, letter *L* and trial period *p* (Fixation, Encoding, Maintenance and Retrieval), we fit the two following models:

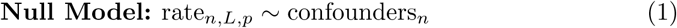

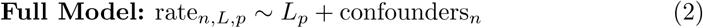

The *L_p_*is a binary time-varying regressor equal to 1 during time bins corresponding to trials in which letter *L* is presented within period *p*, and 0 otherwise. Confounders include trial set size, task period, and spike history. Trial set size (4, 6 or 8) and task period were included as categorical regressors to account for differences in memory load and trial-phase-dependent changes in firing activity. Spike-history confounders were incorporated to control for temporal autocorrelation in neuronal firing. Three history regressors were computed by binning spikes occurring within preceding temporal windows of 11–20, 21–50, and 51–200 bins.

To assess if the *L_p_*regressor improved the model ability to explain the recorded spike trains, we used the information-theoretic measure of bits per spike (BPS) [31], which can be interpreted as a gain in predictability produced by the Full Model compared to the Null model:

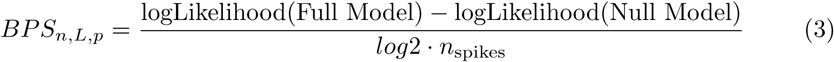

Both models are fit using all data at once. By adding the additional *L_p_*covariate, one necessarily expects the log-likelihood of the full model to increase or remain constant, leading to BPS values superior or equal to zero. To assess whether the BPS increase was significant, we created a surrogate null BPS distribution for every neuronal model to which the empirical BPS was compared. This was done by creating 200 pseudo-trials, randomizing the letter presence across trials. A p-value was then computed from this null distribution and corrected for multiple comparisons (15 = numbers of letter models run for each neuron and trial period) using FDR correction. The models were run using *GLM.jl* library in Julia.

### 4.4 Letter-pair analysis

Similar to the single-letter analysis, we ran, for every neuron, trial period and letter pair, the following models:

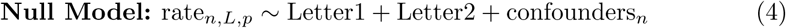

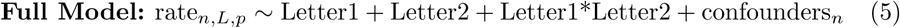

As before, a BPS value was computed for every neuronal model, and a surrogate null distribution obtained by randomizing the letter-pair presence.

For every letter pair and trial period, we computed the maximum-BPS across all 996 neuronal models. To assess the fraction of shared neurons across trial periods driving the maximum-BPS results, we then computed the Spearman’s correlation between the upper-triangular entries of the maximum BPS for all trial periods combinations. To test our two competing hypothesis (conjunctive vs mixed coding), we first found letter-pair selective neurons. For that, we regressed each neuron’s maximum Maintenance BPS across letter pairs onto its maximum letter-pair Fixation BPS, and identified the neurons with residuals greater than two standard deviations above the regression fit as Maintenance-specific compositional encoders. It should be noted that, by experiment and model design, the letter covariates in the model are the same in all stages. To assess if the number of letter-pair encoders found was due to chance, we repeated the analysis 1000 times shuffling the Fixation and Mainteance labels. As a validation, we repeated the analysis for the Encoding period. As a last control in our analysis, we regressed the Fixation BPS onto Maintenance BPS, and following the same steps as above. We then examined whether the letter-pair encoders we identified were located in neurons that also showed strong single-letter selectivity by evaluating the letter-pair BPS, as well as the single-letter BPS for the constituents of the pair. As a control, we also tested if neurons with single-letter selectivity also showed strong letter-pair coding. For that, we took the 14 neurons showing maximum single-letter BPS during the Maintenance period, and evaluated the maximum letter-pair encoding for all possible letter pairs, and for the four trial periods.

## Acknowledgements

We thank the physicians and the staff at Schweizerische Epilepsie-Klinik for their assistance and the patients for their participation.

## Funding

This work was funded by the fellowship B006495 of “la Caixa” Foundation ID 100010434 (E.F.M.), the Swiss National Science Foundation career grant 225979 (T.P.), SNSF 204651 (J.S.) and a UZH Candoc Grant (Grant No. FK-24-025 to F.C.). The funders had no role in the design or analysis of the study.

## Conflict of interest

There is NO competing interests.

## Author contributions

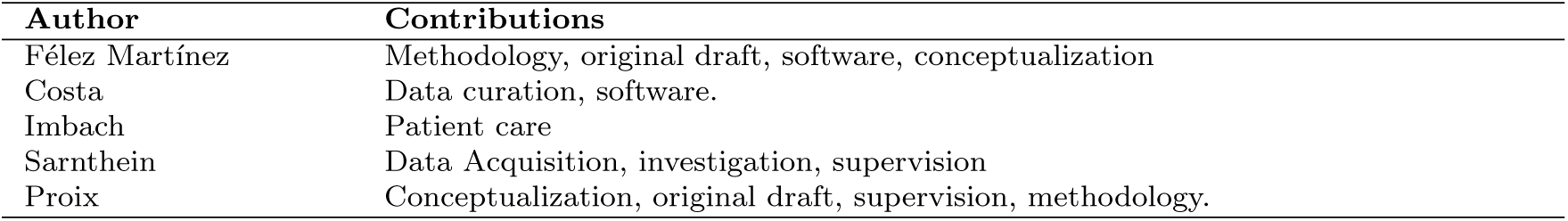

## Appendix A Supplementary information

**Fig. A1:**
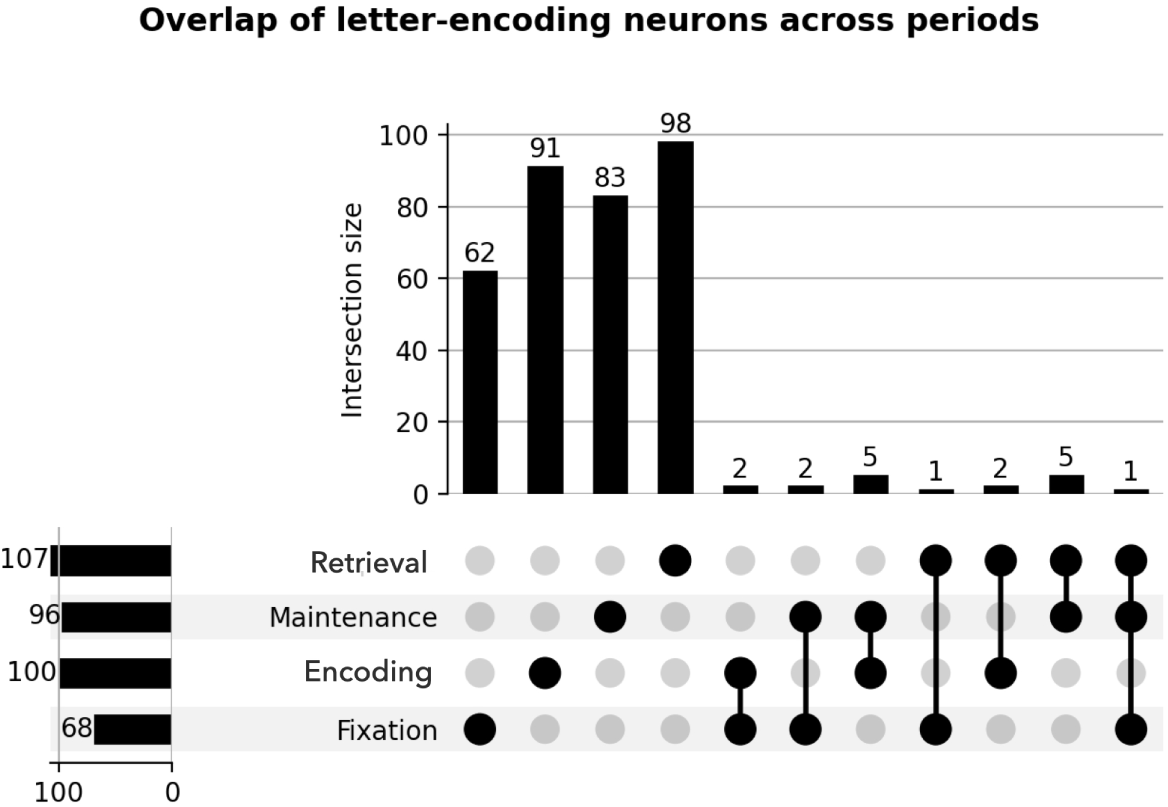
**Distribution of letter-encoding neurons across trial periods,** showing a strong concentration of encoding within specific periods (top left group), with substantially fewer neurons shared across multiple periods (top right group), indicating largely period-specific encoding and limited cross-period overlap.

**Fig. A2:**
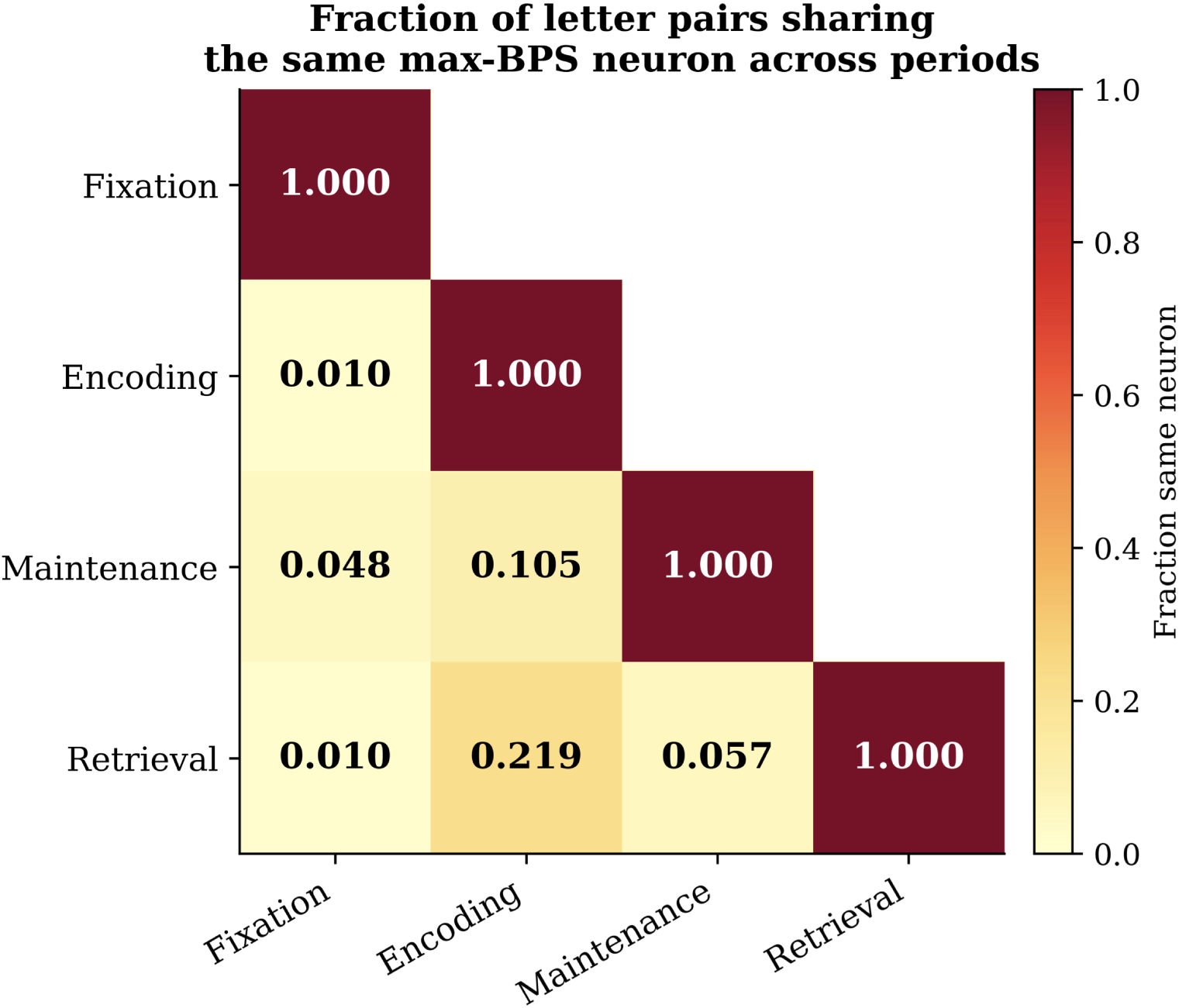
Overlap of max BPS neuron identity across task periods. Each entry shows the fraction of letter pairs for which the same neuron (session, unit) yields the maximum BPS in both periods. Values quantify the stability of the neural code across task periods.

**Fig. A3:**
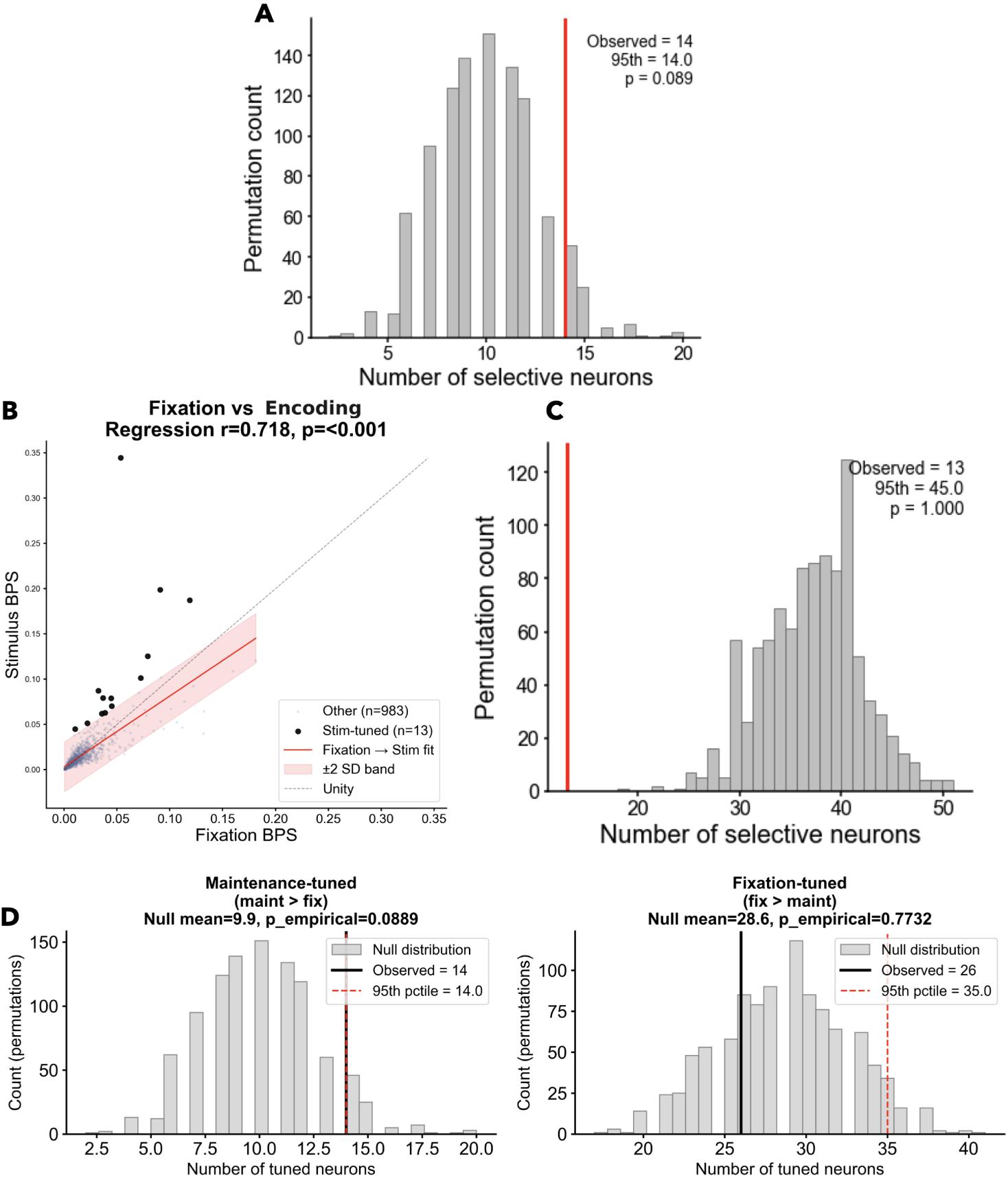
Control analysis reveals the Maintenance-specific effect. **A** We repeated the maintenance-fixation regression analysis 1000 times after shuffling the labels. We found that the 95th percentile of the null distribution was exactly 14 (p = 0.089) **B** We regressed each neuron’s maximum BPS during Encoding period onto its maximum BPS during Fixation, and identified as Maintenance-specific conjunctive coders those neurons whose Maintenance BPS exceeded the regression-predicted value by more than 2 standard deviations of the residual distribution. 14 neurons passed this test, **B** The regression analysis was repeated 1000 times after randomly permuting period labels within each letter pair. The 95th percentile of the resulting null distribution was 45 neurons, yielding an empirical p-value of 1. **C** We did the same analysis as above (left panel, Fig. 5) but now inverting the regression (i.e., regressing Fixation BPS to Maintenance BPS). We found 26 neurons, but the permutation test yields p = 0.77 (right panel).

**Fig. A4:**
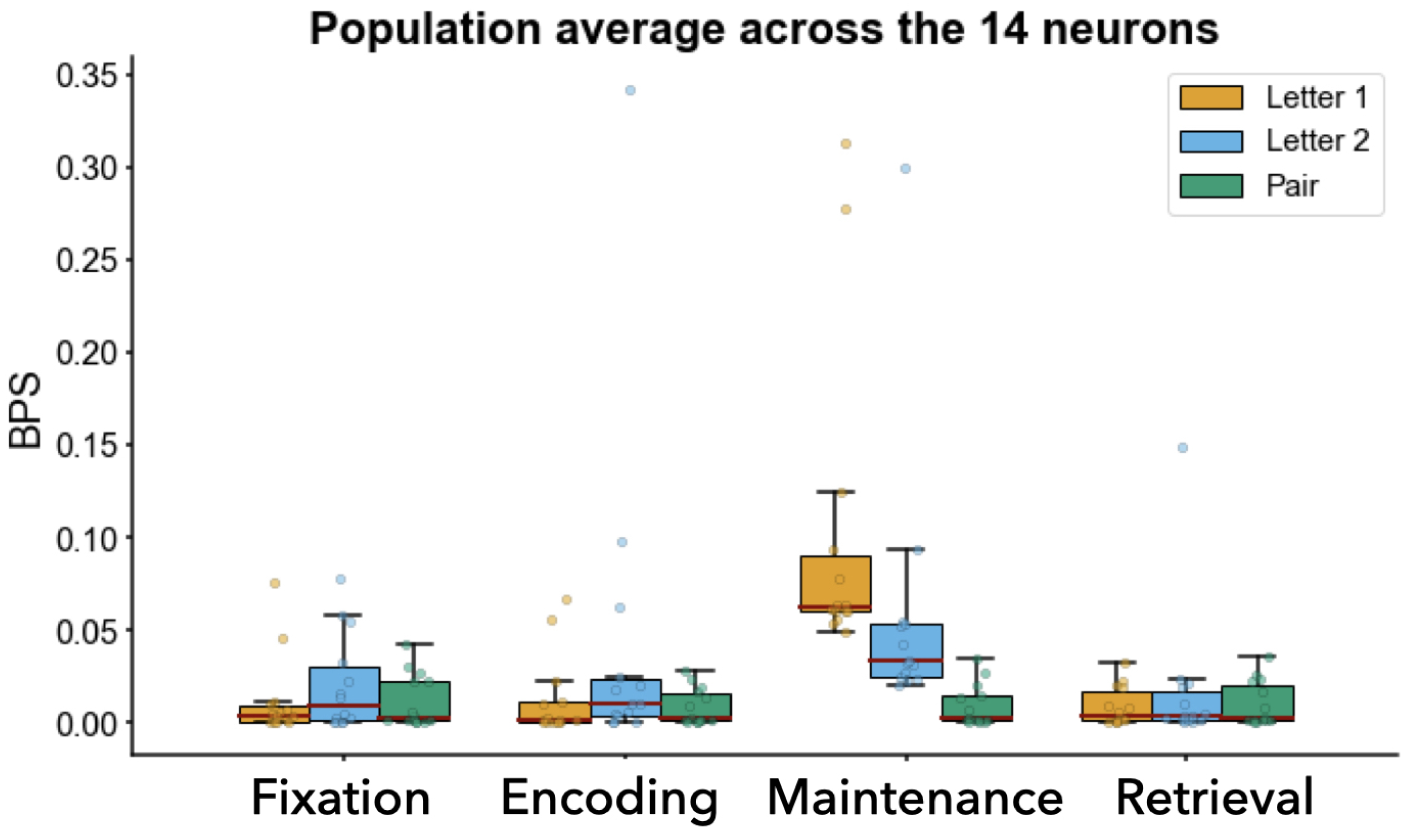
Control analysis: testing whether pair selectivity arises from strong single-letter tuning. Neurons were selected independently of any pair analysis by ranking all recorded units according to their maximal single-letter BPS during the Maintenance period (14 were selected). For each neuron, BPS from the single-letter analysis were obtained for (i) the most selective letter (orange), (ii) the second-most selective letter (blue), and (iii) the strongest letter pair response (green; maximum across all possible pairs for that neuron). Despite strong tuning to individual letter, pair responses remain low across periods, demonstrating that pair selectivity does not trivially emerge from single-letter encoding.

## Notes

### Competing Interest Statement

The authors have declared no competing interest.

